# TNF signaling is required for castration-induced vascular damage preceding prostate cancer regression

**DOI:** 10.1101/2022.02.05.479251

**Authors:** John J. Krolewski, Shalini Singh, Kai Sha, Neha Jaiswal Agrawal, Steven G. Turowski, Chunliu Pan, Laurie J. Rich, Mukund Seshadri, Kent L. Nastiuk

**Affiliations:** Department of Cancer Genetics & Genomics; Roswell Park Comprehensive Cancer Center, Buffalo, NY 14263; Department of Cell Stress Biology; Roswell Park Comprehensive Cancer Center, Buffalo, NY 14263; Laboratory of Translational Imaging, Center for Oral Oncology; Roswell Park Comprehensive Cancer Center, Buffalo, NY 14263; Department of Urology; Roswell Park Comprehensive Cancer Center, Buffalo, NY 14263

**Keywords:** photoacoustic imaging, contrast-enhanced ultrasound, power Doppler, endothelial, sTNFR2-Fc, cancer therapy, mouse models

## Abstract

The mainstay treatment for locally advanced, recurrent, or metastatic prostate cancer (PrCa) is androgen deprivation therapy (ADT). ADT causes prostate cancers to shrink in volume, or regress, by inducing epithelial tumor cell apoptosis. In normal, non-neoplastic murine prostate, androgen deprivation via castration induces prostate gland regression that is dependent on TNF signaling. Besides this direct mechanism of action, castration has also been implicated in an indirect mechanism of prostate epithelial cell death which has been described as vascular regression. The initiating event is endothelial cell apoptosis and/or increased vascular permeability. This subsequently leads to reduced blood flow and perfusion, and then hypoxia, which may enhance epithelial cell apoptosis. Castration-induced vascular regression has been observed in both normal and neoplastic prostate. We used photoacoustic, power Doppler, and contrast-enhanced ultrasound imaging, and CD31 immunohistochemical staining of the microvasculature to assess vascular integrity in the period immediately following castration, enabling us to test the role of TNF signaling in vascular regression. In two mouse models of androgen-responsive prostate cancer, TNF signaling blockade using a soluble TNFR2 ligand trap reversed the functional aspects of vascular regression as well as structural changes in the microvasculature, including reduced vessel wall thickness, cross-sectional area and vessel perimeter length. These results demonstrate that TNF signaling is required for vascular regression, most likely inducing endothelial cell apoptosis and increasing vessel permeability. Since TNF is also the critical death receptor ligand for prostate epithelial cells, we propose that TNF is a multi-purpose, comprehensive signal within the prostate cancer micro-environment mediating prostate cancer regression following androgen deprivation.

**SIGNIFICANCE:** These studies define TNF as the mediator of androgen deprivation therapy-induced functional and structural vascular damage in prostate tumors.

## Introduction

Prostate cancer (PrCa) is the second leading cause of cancer-related mortality for US men (1). While localized PrCa can be cured in many men by surgery or radiation, androgen deprivation therapy (ADT) is the mainstay treatment for locally advanced, recurrent, or metastatic PrCa (2). Prostate cancers regress following ADT due to the death of epithelial-derived tumor cells. Indeed, one of the earliest morphological descriptions of apoptotic cell death was in the luminal epithelial cells of the rat prostate gland following castration (3). Tissue reconstitution experiments by Cunha and colleagues demonstrated that androgen blockade causes prostate cancer regression indirectly via stromal-derived soluble factors that act on the tumor epithelium (4). For example, we previously demonstrated that stromal-derived tumor necrosis factor (TNF) – the prototypical death receptor ligand – mediates castration-induced regression in normal murine prostates (5). Specifically, castration-induced regression of the normal prostate is reduced in both Tnfr1 and Tnf deleted mice as well as mice treated with the TNF ligand trap drug etanercept (5).

Although castration induced TNF signaling directly triggers epithelial cell apoptotic death in prostate cancers, there is substantial evidence that androgen withdrawal also affects the prostate tumor microvasculature (6; 7). Specifically, in rodent models, castration leads to an early induction of endothelial cell apoptosis and micro-vessel leakiness (8), a reduction in blood flow (9; 10) and subsequent reduced perfusion of tumor tissue (11) which leads to ischemia and acute hypoxia (12-15). A similar process of castration induced microvasculature damage, caused by paracrine mediators, was observed in primary human xenografts (7). Therefore, it has been proposed that there is a vascular phase or component to castration-induced regression of prostate cancers during which damage (apoptosis, leakiness) to the tumor microvasculature eventually produces tumor hypoxia. We suggest that the hypoxic state then either initiates or accelerates necrotic and/or apoptotic death of the epithelial tumor cells. The signaling mechanism that mediates ADT-induced vascular damage is unknown, but TNF has been shown to induce both endothelial cell apoptosis and vascular permeability (16; 17) and is therefore a candidate for a secreted protein that can function in a paracrine manner to trigger vascular regression.

To test our hypothesis that TNF mediates the vascular component of ADT-induced regression, we monitored the effects of castration on the functional and structural properties of the tumor microvasculature in a c-Myc driven, androgen-sensitive prostate cancer model. We report that castration induced micro-vessel structural damage, blood flow, perfusion, and oxygenation in the tumor, confirming and extending prior descriptions of the vascular events accompanying regression in prostate cancers. All of these functional and structural vascular alterations induced by castration were reversed by concurrent TNF signaling blockade, demonstrating that endogenous TNF signaling is necessary for castration-induced vascular regression and likely acts at the initiating step of the process, mediating endothelial cell apoptosis and vascular permeability. Combined with our previous studies, our findings suggest a comprehensive role for TNF signaling in the ADT-induced regression of prostate cancers.

## Materials and Methods

### Cell culture

The Myc-CaP cell line was established from the Hi-Myc transgenic mouse prostate (18) stably transfected with firefly luciferase under the androgen response element promoter (19). Myc-CaP/ARE-luc cells were stored as low-passage aliquots, tested for mycoplasma and pathogens, and periodically renewed from the frozen stocks. Cells were grown in Dulbecco’s Modified Eagle Medium containing glutamine and antibiotics with 10% fetal calf serum.

### Animals

All animal studies were performed in accordance with the National Institute of Health Guidelines for the Care and Use of Laboratory Animals and approved by the Roswell Park Institutional Animal Care and Use Committee (#1308M). Experimental studies were performed using two mouse models of PrCa, subcutaneous allografts of Myc-CaP cells and an autochthonous genetically engineered mouse model, prostate-specific Pten deletion (20). All mice were maintained on a 12-hour light and 12-hour dark cycle and had regular chow ad libitum.

To establish the Myc-CaP tumors, both flanks of male FVB/NCr mice (8-12 week old, Charles River Labs) were injected with 5×10^5^ Myc-CaP/ARE-luc cells re-suspended in equal volumes of RPMI media and Matrigel (Corning, Bedford, MA). Tumor development was monitored by palpation and when evident, quantitated by high-frequency ultrasound (HFUS) imaging, as previously described (21). Mice were treated and castrated or sham castrated within 17 days of tumor cell injection. Average tumor volume at the time of castration was 382 +/- 120 mm^3^.

To produce prostate-specific PTEN loss induced PrCa-bearing mice (20), male transgenic mice expressing probasin driven Cre recombinase (PB-cre4; obtained from NCI) were crossed with female floxed Pten mouse (Pten^LoxP/LoxP^; obtained from JAX). The resulting pups were individually identified by implantation of a p-Chip (Pharmaseq, Monmouth Junction, NJ) and genotyped for the Cre transgene and the LoxP containing Pten alleles. At puberty, the probasin promoter is activated in prostate secretory epithelium, to cause epithelial cell specific deletion of Pten and thereby loss of PTEN activity. Prostate adenocarcinomas form with complete penetrance between 3 and 7 months. The animals were therefore monitored for tumor development using HFUS imaging beginning at 12 weeks. Tumor-bearing animals were enrolled for experimental manipulation when tumor was between 300 mm^3^ and 700 mm^3^ (mean 436+/-31 mm^3^).

TNF signaling was blocked by treating mice with etanercept (Amgen), using a regimen previously demonstrated to block castration-induced regression of the normal prostate gland (5). Etanercept is the soluble extracellular portion of TNFR2 coupled to the IgG Fc domain (sTNFR2-Fc). At three and one day prior to castration, tumor-bearing mice were treated with sTNFR2-Fc by intraperitoneal (ip) injection at 4 mg/kg. Soluble TNF (PeproTech, Rocky Hill, NJ) was injected (27 µg/kg) ip at the time of castration, at a dose sufficient to restore castration-induced prostate regression in TNF-deficient mice (5). Control mice received PBS on the same schedule. Mice were surgically castrated on day zero as previously described (22). Briefly, animals were anesthetized using 2.5% Isoflurane (Benson Medical Industries, Markham, ON, Canada), and maintained on 1% Isoflurane in oxygen. Testes were removed via bilateral scrotal incision, the blood vessels and vas deferens ligated, and scrotal incisions closed by suturing (rather than stapling) to enable imaging of the peritoneum.

### Anatomic and functional imaging

Ultrasound imaging with co-registered photoacoustic imaging (PAI), power Doppler, contrast-enhanced ultrasound (CE-US) and B-mode high-resolution HFUS imaging were performed using a 256 element, 21MHz linear-array transducer (LZ-250) and the Vevo LAZR system (VisualSonics Inc., Toronto ON, Canada). After mice were anesthetized and depilated, B-mode ultrasound images were acquired from the peritoneum, and 3D reconstructions of the prostate tumors were computed using Amira 3D visualization software (FEI Visualization Sciences Group) (21). For power Doppler sonography parameters used for acquisition were: operating frequency: 16MHz, pulse repetition frequency: 2kHz, Doppler gain: 40, depth: 20.00 mm, width: 23.04 mm, clutter/wall filter: medium. To enable accurate comparison of power Doppler data, the relative change in power Doppler signal was calculated for the 3D ROI covering the entire tumor. Multispectral PAI was performed to obtain measurements of oxygen saturation (sO2). PAI parameters used were: Operating frequency 21 MHz, Depth: 23.00 mm, Width: 23.04 mm, Wavelength: 750/850 nm, total hemoglobin concentration threshold (Hbt) was 20 arbitrary units, acquisition mode: sO2/Hbt. The photoacoustic gain was kept at 43 dB and dynamic range at 20 dB for all studies. PAI based measurements of oxygen saturation were calculated using the two-wavelength approach (750/850 nm) as described (23; 24). Nonlinear Contrast Mode imaging was performed to detect the presence of Vevo MicroMarker contrast agent (VisualSonics, Toronto, ON, Canada). The contrast agent consists of phospholipid shell microbubbles filled with nitrogen and perfluorobutane (2.3 to 2.9 μm in diameter). A bolus injection of the contrast agent (1×10^8^ microbubbles) was administered via tail vein injection using a 25-gauge needle. Images were acquired using the following parameters: Operating frequency 18MHz, Depth: 20.00 mm, Width: 23.04 mm, with 35dB contrast gain, gate size 6. Nonlinear detection of the contrast signal was done in 3D by moving the transducer through the volume of the tumor at a step-size of 0.152 mm. Multimodal imaging datasets were processed offline using VEVO CQ software utilizing manually drawn tumor regions with perfusion parameters derived from intratumoral signal intensity time curves. All imaging datasets were analyzed using Vevo LAB (v.1.7.2) workstation software.

### Immunohistochemistry

Mice were sacrificed 24h after castration and tumors collected for histology. Tumors were fixed in formalin-free immunohistochemistry zinc fixative (BD Pharmingen, San Diego, CA) and 4 µm paraffin sections obtained. Vessels were identified by immunohistochemical staining for CD31 (rat anti-mouse CD31, clone MEC13.3, #550274, BD Pharmingen) using an autostainer (Agilent/DAKO Carpinteria, CA) as reported (11). Images were digitized using the ScanScope XT system. The entire tumor area was delineated using Aperio ImageScope v11 and vasculature integrity was analyzed using the default parameters of the automated micro-vessel analysis algorithm v1.1 (Aperio Technologies, Vista, CA).

### Statistical analysis

Tumor volume and ELISA TNF protein levels were compared using one-way ANOVA and Dunnett’s test post-hoc. Saturated oxygenation, vascularity, and perfusion were analyzed using two-way ANOVA employing Tukey’s honestly significant difference test post-hoc to compare treatments over time. ANOVA two-tailed p values <0.05 were considered significant, and, if reached, post-hoc testing was performed. Unpaired Student’s t-test were employed for single-treatment comparisons of histological parameters, and paired Student’s t-test for single-treatment analyzed using pre- and post-imaging. Two-tailed p values <0.05 were considered significant. All statistical analyses were performed using JMP Pro 11.0 software (SAS).

### Data and materials availability

The data generated in this study are available within the article and its supplementary data files.

## Results

To determine if TNF signaling plays a role in the vascular events accompanying castration-induced regression of prostate cancer, we employed a subcutaneously implantable Myc-CaP tumor model. The cell line used in this allograft model is derived from a prostate cancer that developed in Hi-MYC mice (18; 25), which express c-MYC oncogene in the prostate under the control of an androgen-regulated promoter (ARR_2_/probasin-*Myc*). Since c-MYC amplification or over-expression is frequently seen in human prostate cancers (26), this model is relevant to human prostate cancers. We recently studied castration-induced vascular changes in Myc-CaP allografts implanted in mice, as part of a study to evaluate vascular targeting as a therapeutic strategy for treating prostate cancer (27). In that report, we documented some of the vascular changes that are have been previously described in castration induced vascular regression (6). In particular, one day following castration, we observed a reduction in tissue perfusion. Importantly, the effect was similar in tumors implanted subcutaneously and orthotopically, allowing us to reliably use the subcutaneous model in the present studies, since signals from some of the tissue imaging modalities we employ to assess vascular function are attenuated by tissue, making it difficult to make measurements on internal organs such as the prostate.

Myc-CaP tumors were implanted subcutaneously and tumor-bearing mice imaged by high frequency ultrasound (HFUS) to monitor tumor size. Once tumors reach about 350-400 mm^3^, mice were castrated and one day later (before there is a significant reduction in tumor volume, Supplementary Figure 1A), we measured changes for each step of the ordered sequence of the alterations that define the vascular contribution to tumor regression: i) microvasculature damage; ii) reduced blood flow; iii) reduced tissue perfusion; and iv) tissue hypoxia. To specifically test the role of TNF signaling, experiments were performed in the presence or absence of sTNFR2-Fc – a soluble form of the TNFR2 extracellular domain fused to the immunoglobulin Fc protein – that binds TNF and prevents signaling in the target cell.

### TNF is necessary for castration-induced microvasculature damage in Myc-CaP allografts

To test the role of TNF signaling in vessel structural changes, mice bearing Myc-CaP tumors were castrated and tumor was excised and formalin-fixed one day post-castration from both vehicle and sTNFR2-Fc treated animals. The microvasculature network was visualized with CD31 immuno-histochemical staining of tissue sections (Fig. 1A) and then analyzed using an automated algorithm that determines micro-vessel density and related parameters. Treatment with sTNFR2-Fc increased micro-vessel density in two of four tumors from mice (Fig. 1B). In addition, intratumoral area occupied by vessels, the vessel perimeter and the thickness of the vessel walls were all increased by sTNFR2-Fc treatment in all four tumors examined (Fig. 1C-E). We previously reported castration decreased the number of vessels in both subcutaneous and orthotopic Myc-CaP allografts by ∼30% after one day (11). Taken together, these data are consistent with TNF mediating the vascular structural damage induced by castration in prostate tumors.

**Figure 1.**
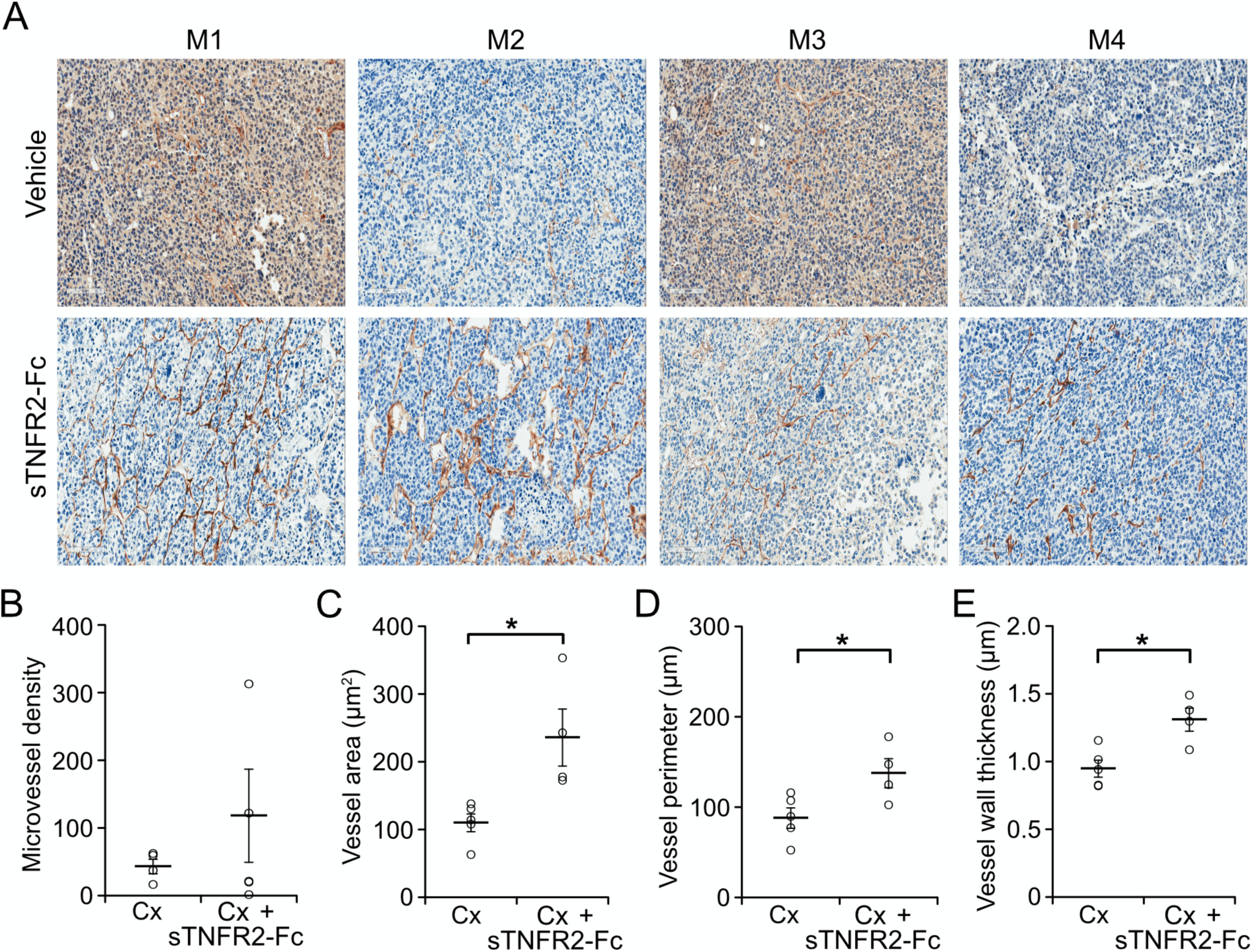
Castration-induced vascular damage is reversed by TNF signaling blockade. **A**, Representative sections of CD31 immunoreactivity in Myc-CaP prostate tumor from four mice (M1-M4) castrated (upper panels) or castrated and treated with sTNFR2-Fc treated (lower panels). **B**, Micro-vessel density (vessels per square millimeter). **C**, Vessel cross-sectional area, in µm^2^; **D**, Vessel perimeter length, in µm; **E**, Vessel wall thickness, in µm. Columns, mean of five tumors from vehicle-treated and four sTNFR2-Fc-treated castrated mice, bars = SEM.

### Castration-induced reduction in blood flow and perfusion is prevented by blocking TNF signaling

Next, we used power Doppler imaging to quantitate the intensity of intratumoral blood flow. Doppler imaging uses high-frequency sound waves to visualize blood flow magnitude (28). Power Doppler imaging has increased sensitivity versus color Doppler imaging, allowing more complete vessel function imaging by integrating speed and directional information (29; 30). Castration reduced blood flow inside the tumors (Fig. 2A, 2B) but blood flow was not changed when TNF signaling was blocked using sTNFR2-Fc (Fig. 2C, 2D). The castration-induced decrease in overall tumor blood flow was TNF-dependent one day after castration, and while blood flow continued to decline, sTNFR2 treatment did not affect the castration-induced change in blood flow after four days castration (Fig. 2E, individual tumor changes at one day after castration are shown in Fig. 2F). Serial measurement of the change in blood flow inside the same tumor confirmed the castration reduction of blood flow was acutely TNF dependent (Fig. 2G).

**Figure 2.**
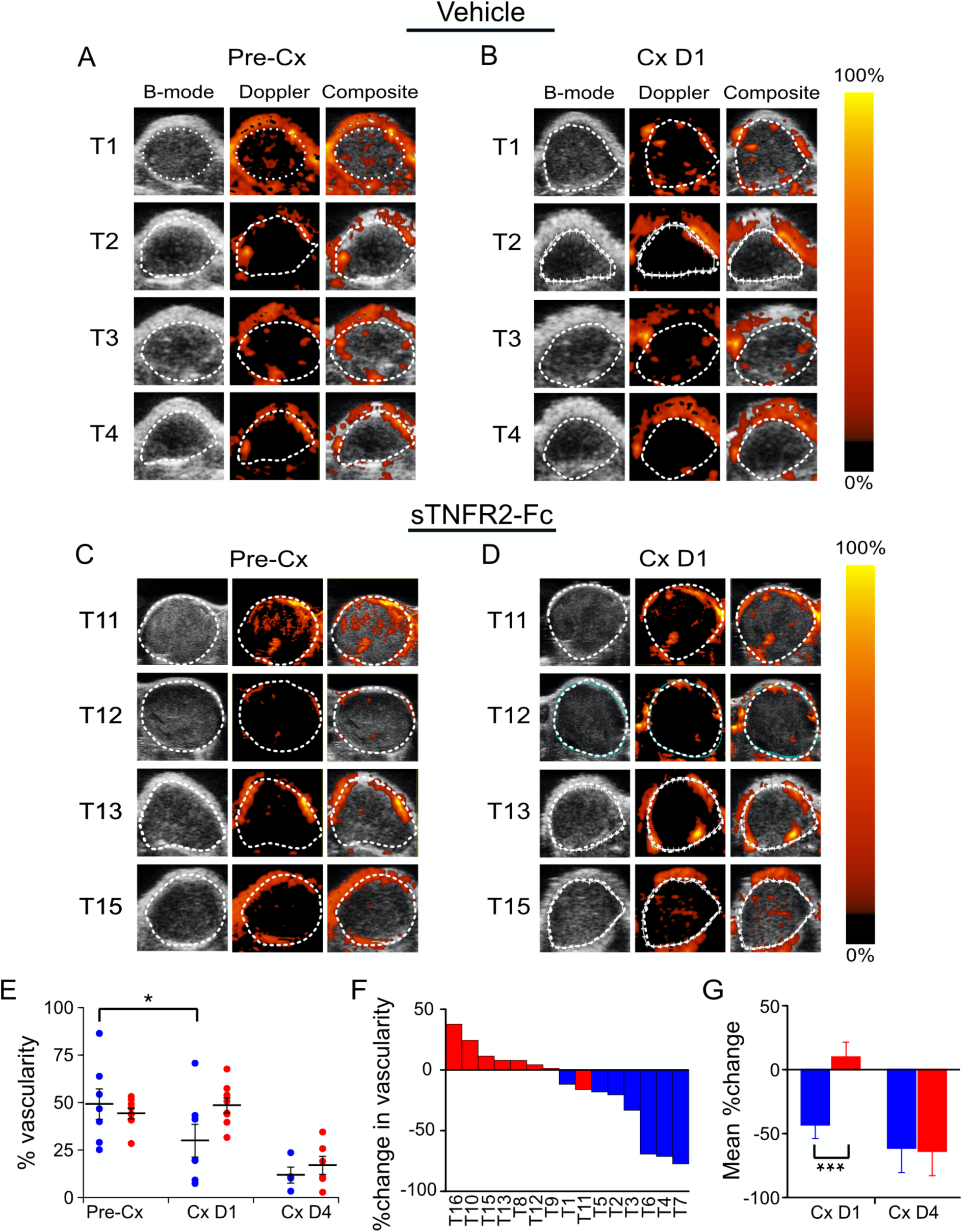
Castration reduction of intratumoral blood flow in Myc-CaP tumor was dependent on TNF signaling. **A**, Power Doppler (PD) images of subcutaneous Myc-CaP tumors pre-castration (Pre-Cx) of mice treated with PBS (Vehicle), tumors 1-4 (of 8 evaluable tumors). Left to right: Gray-scale ultrasound image (B-mode); PD pseudo-colored image to illustrate blood flow level (Doppler); Composite of PD image overlaid on the B-mode image (Composite). **B** PD images of tumors in panel A, one day after castration (Cx D1). **C**, PD images of a second set of subcutaneous Myc-CaP tumors pre-castration of mice treated with sTNFR2-Fc, tumors 11, 12, 13, 15 (of 8 evaluable tumors). **D**, PD images of tumors in panel C, one day after castration. **E**, Mean PD signal (% vascularity) pre-castration, and at one and four days after castration from tumors in vehicle treated (blue, D1 n=8, D4 n=7) and sTNFR2-Fc treated mice (red, D1, D4 n=8). **F**, Waterfall plot of % change in vascularity in individual tumors at one day after castration. **G**, Mean % change in paired measures of vascularity pre-castration versus D1 or D4 after castration. Columns are means and bars are SEM, *p<0.05, ***p<0.001.

To test if the changes in blood flow also resulted in reduced tumor perfusion, Myc-CaP allografts were evaluated using contrast-enhanced ultrasound (CE-US) prior to and one-day after castration of the host animal. Relative perfusion was determined using contrast agent accumulation in tumor capillaries and quantitated from a maximum intensity projection based on contrast accumulation. Figure 3 shows castration reduced intratumoral perfusion (Fig. 3A, 3B). However, castration did not reduce intratumoral perfusion in sTNFR2-Fc pre-treated host mice (Fig. 3C, 3D). Tumors in sTNFR2-Fc pre-treated host mice were highly perfused, and this was also not changed by castration (Fig. 3E). Castration-reduction of intratumoral perfusion was dependent on TNF signaling (individual tumor changes shown in Fig. 3F, mean changes shown in Fig. 3G).

**Figure 3.**
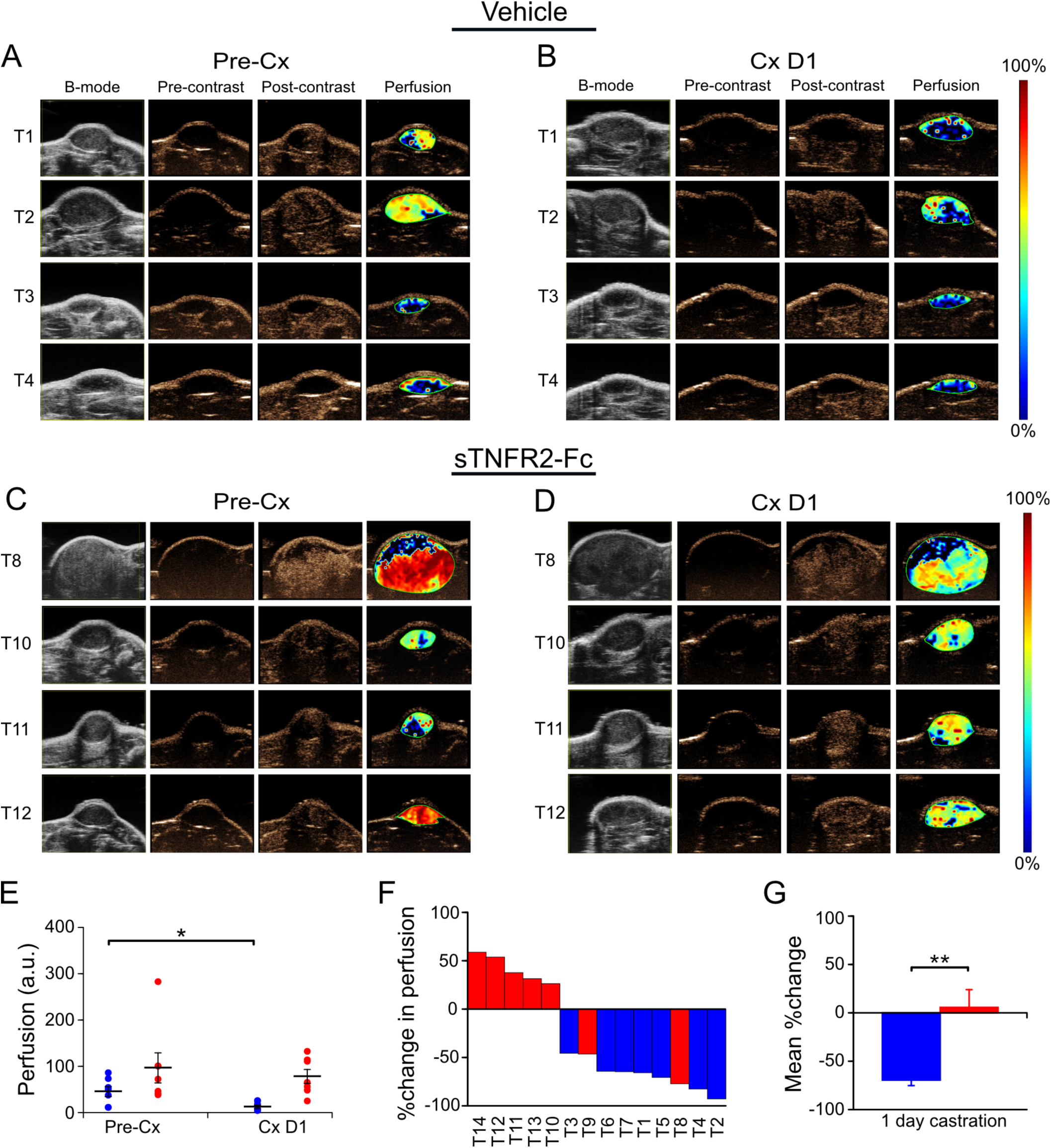
TNF signaling was necessary for castration-induced reduction in perfusion of Myc-CaP tumor. **A**, Contrast enhanced-ultrasound (CE-US) images of subcutaneous Myc-CaP tumors pre-castration (Pre-Cx) of mice treated with PBS (Vehicle), tumors 1-4 (of 7 evaluable tumors) are shown. Left to right: Gray-scale ultrasound image (B-mode); contrast-mode image prior to contrast agent injection (Pre-contrast); contrast-mode image after injection at the peak enhancement of contrast (Post-contrast); pseudo-colored image of the change in contrast enhancement (Perfusion). **B**, CE-US images of tumors 1-4 in A after one day castration (Cx D1). **C**, CE-US images of a second set of subcutaneous Myc-CaP tumors pre-castration of mice treated with sTNFR2-Fc, tumors 8, 10, 11, 12 (of 7 evaluable tumors) are shown. **D**, CE-US images of tumors in panel C, one day after castration, **E**, Mean perfusion pre-castration and post-castration in tumors in vehicle treated (blue) and sTNFR2-Fc (red) treated mice. **F**, Waterfall plot of %change in perfusion in individual tumors (columns). **G**, Average %change in perfusion pre-castration and post-castration. E, G: Columns are means and bars are SEM, *p<0.05, **p<0.01.

### TNF is required for castration-induced hypoxia in Myc-CaP allografts

Given our observations of reduced blood flow and perfusion, we next sought to determine the level of tissue oxygenation, using photoacoustic imaging (PAI) to measure oxygen saturation. PAI can discriminate the absorption spectra of endogenous oxy-hemoglobin from deoxy-hemoglobin, enabling real-time 3D visualization of microvasculature and quantitation of changes in the percent hemoglobin oxygen saturation (%sO_2_) (31). As in prior experiments, Myc-CaP allograft volumes were quantitated using HFUS imaging and tumor-bearing mice were pre-treated with sTNFR2-Fc (or vehicle) to block TNF signaling, and then castrated. One day after castration, photoacoustic signal was decreased (Fig. 4A) but no castration-induced change in photoacoustic signal was seen if TNF signaling was blocked using sTNFR2-Fc (Fig. 4B). Castration induced a relative decrease in intratumoral hemoglobin oxygen saturation (%sO_2_) in one day after castration, but not when TNF signaling was blocked in host mice using sTNFR2-Fc (Fig. 4C, individual tumor changes shown Fig. 4D). The castration-induced hypoxia was not detectable by four days after castration. Total intratumoral hemoglobin was not changed by either castration or sTNFR2-Fc treatment (Fig. 4E). Comparison of paired oxygenation changes in the same tumor revealed the castration induction of hypoxia was TNF dependent (Fig. 4F).

**Figure 4.**
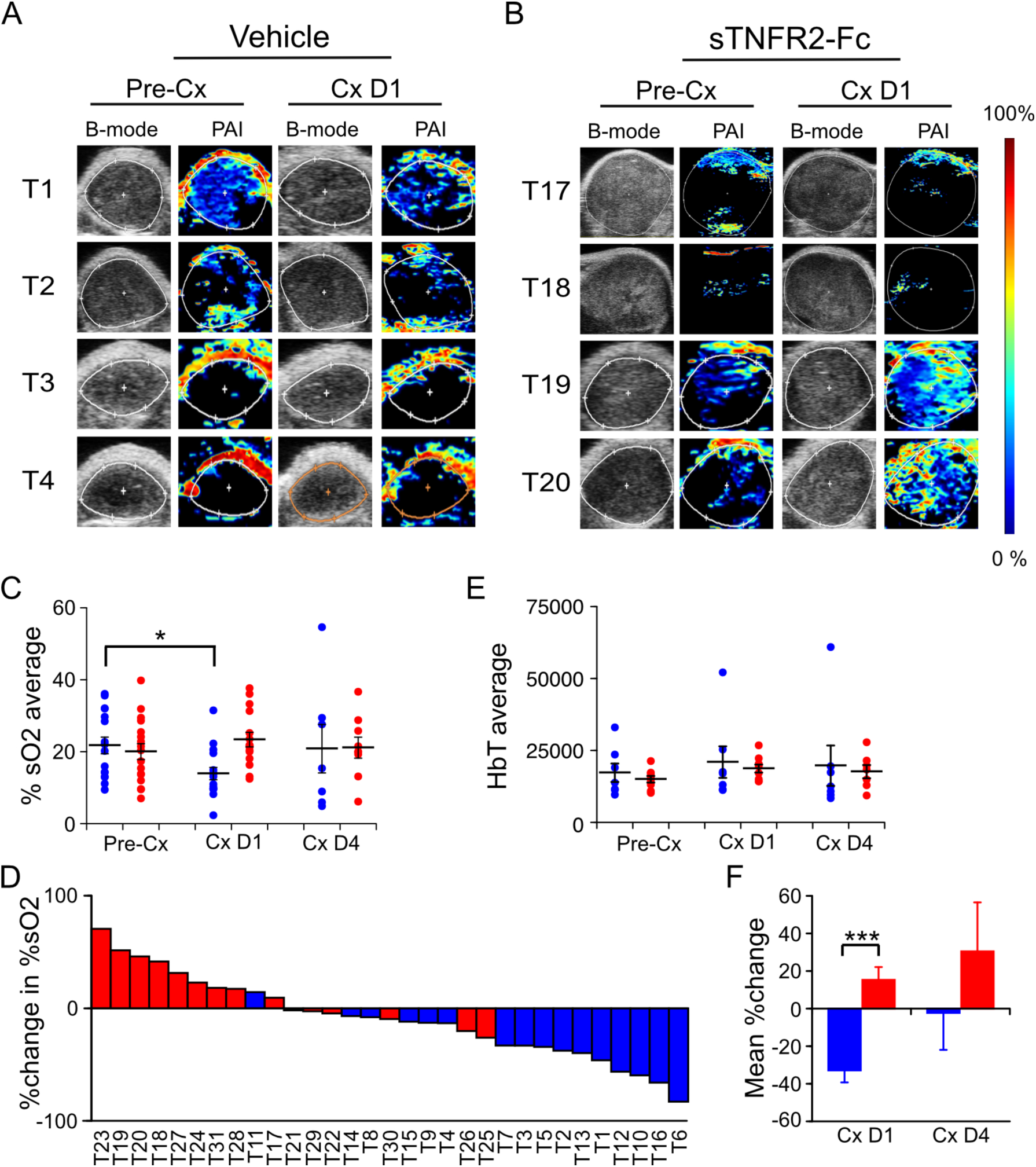
Castration induced hypoxia in Myc-CaP tumor is reversed by TNF signaling blockade. **A** and **B**, Photoacoustic images (PAI, pseudo-colored) and ultrasound (B-mode) of Myc-CaP subcutaneous tumor of four tumors from each group, pre and post castration from vehicle treated mice (**A**), or sTNFR2-Fc treated mice (**B**). **C**, Mean PAI signal (%sO_2_) from tumors pre-castration (n=16), and at one (n=15), and at four (n=9) days post-castration in vehicle treated (blue) or in sTNFR2-Fc treated (red) mice. **D**, Mean total hemoglobin from tumors pre-castration (n=16), and at one (n=16), and at four (n=14) days post-castration in vehicle treated (blue) or in sTNFR2-Fc treated (red) mice. **E**, Waterfall plot of change in %sO_2_ in individual tumors at one day post-castration versus pre-Cx. **F**, Mean change in paired measures of %sO_2_ pre-castration versus D1 or D4 post-castration. Mean (columns) and SEM (bars). *p<0.05, ***p<0.001.

### TNF is required for castration-induced hypoxia in an autochthonous prostate cancer model

To determine whether castration induces TNF-dependent vascular damage in a more clinically relevant prostate cancer model, we examined vascular change after castration in the endogenously arising PrCa tumors in PbCre4 x Pten^fl/fl^ mice (20). Tumorigenesis in this model is driven by Pten gene loss in the prostate epithelium. PTEN loss – similar to c-MYC gain of function – is one of the most frequent genetic lesions in both localized and metastatic human PrCa (32) and PTEN loss predicts outcome in patients (33). Like human PrCa, these autochthonous tumors grow slowly and regress after castration (20). We first tested whether TNF signaling acutely affected castration-induced regression or TNF expression in this PrCa model. When tumors developed to between 300 mm^3^ and 500 mm^3^ (mean 347 +/- 28 mm^3^), mice were pre-treated with sTNFR2-Fc or vehicle, and castrated.

We tested if castration induced hypoxia in this model, and whether hypoxia was dependent on TNF signaling. When PrCa tumor volume was 300 mm^3^ - 700 mm^3^, mice were treated with sTNFR2-Fc and castrated. Tumor hypoxia levels were then assessed serially using PAI. Castration induced a decrease of %sO_2_ in the tumors of vehicle treated mice after one day (Fig. 5A, 5C), but not after four days (Fig.5C). This castration induced reduction in %sO_2_ was abrogated in the tumors of mice pre-treated with sTNFR2-Fc (Fig. 5B, 5C). The level of absolute PA signal (∼4% sO_2_) in this PrCa model with black pigmentation was much lower than in the Myc-CaP allografts (∼20% sO_2_) in white colored hosts, and in our previous report for a subcutaneous tumor model (∼40% sO_2_) in albino hosts (24). Despite the reduced absolute signal strength, intratumoral paired change in %sO_2_ was uniformly decreased in vehicle-treated mice one and four days after castration (blue, Figs. 5D, 5E). This effect was blocked in most tumors of sTNFR2-Fc pre-treated mice at both one and four days after castration (red, Figs. 5D, 5E). The average castration-induced a 50% decrease %sO_2_ (hypoxia) was reversed by sTNFr2-Fc pre-treatment (Fig. 5F). Power Doppler based measurements of blood flow and CE-US measurement of tumor perfusion were unchanged one or four days after castration in this PrCa model (data not shown).

### TNF is not sufficient for castration induced hypoxia in Myc-CaP allografts

**Figure 5.**
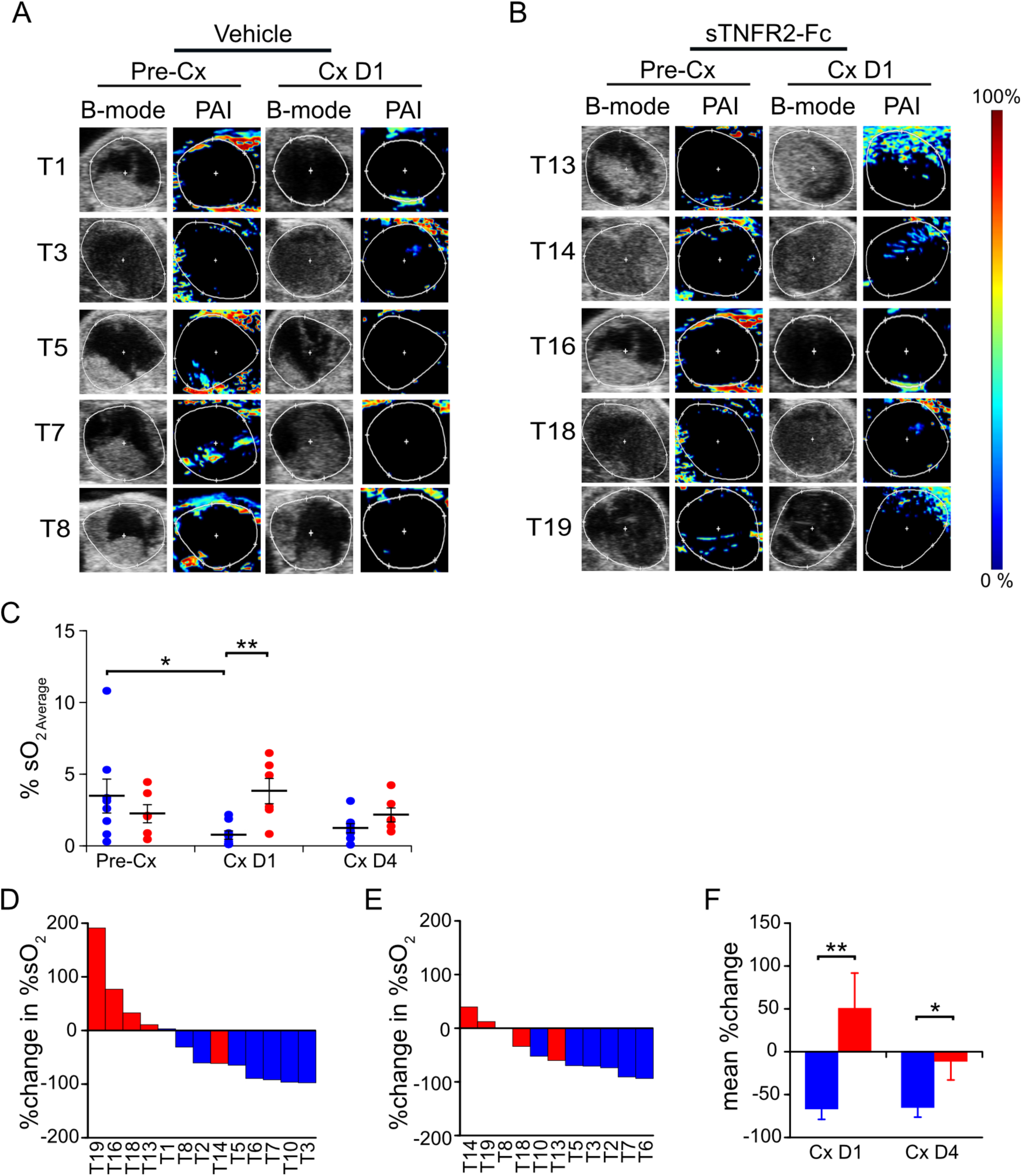
Castration induced hypoxia is reversed by TNF blockade in prostate tumors of PbCre4 x Pten^fl/fl^ mice. **A, B**, Photoacoustic images (PAI, pseudo-colored) and ultrasound (B-mode) of five tumors pre and one day post castration from vehicle treated mice (A), or sTNFR2-Fc treated mice (B). **C**, Intra-tumoral mean PAI intensity (%sO_2 Average_) pre-castration, and at one and at four days post-castration in vehicle treated (n = 8, 8, 7 respectively, blue) or in sTNFR2-Fc treated (n = 6, 5, 4 respectively, red) mice. **D**, Waterfall plot of change in intra-tumoral %sO_2_ one day after castration in mice treated with sTNFR2-Fc (red) or vehicle (blue). **E**, Waterfall plot of change in intra-tumoral %sO_2_ four days after castration in mice treated with sTNFR2-Fc (red) or vehicle (blue). **F**, Change in paired measures of intra-tumoral %sO_2_ pre-castration versus D1 or D4 after castration. Mean (columns or lines) and SEM (bars). *p<0.05, **p<0.01, ***p<0.001.

Finally, since blocking TNF signaling was sufficient to abrogate castration-induced hypoxia, we tested whether soluble TNF was sufficient to induce hypoxia in Myc-CaP tumor. When Myc-CaP allografts became palpable, tumor volume was quantitated, PAI performed and then host mice were treated with TNF. TNF was administered at a dose sufficient to rescue prostate regression in TNF deficient mice (5). Surprisingly, tumor oxygenation increased one day after TNF injection (Fig. 6A). TNF-injection induced change of %sO2 in individual tumors ranged from 20% to 240% (Fig. 6B). Moreover, TNF protein levels were unchanged in soluble lysates of whole tumor after castration (Supplementary Fig. S1B). These data suggest that in the subcutaneous Myc-CaP model, TNF signaling is not triggered by a castration induced increase in TNF protein, but instead involves modulation of downstream signaling.

**Figure 6.**
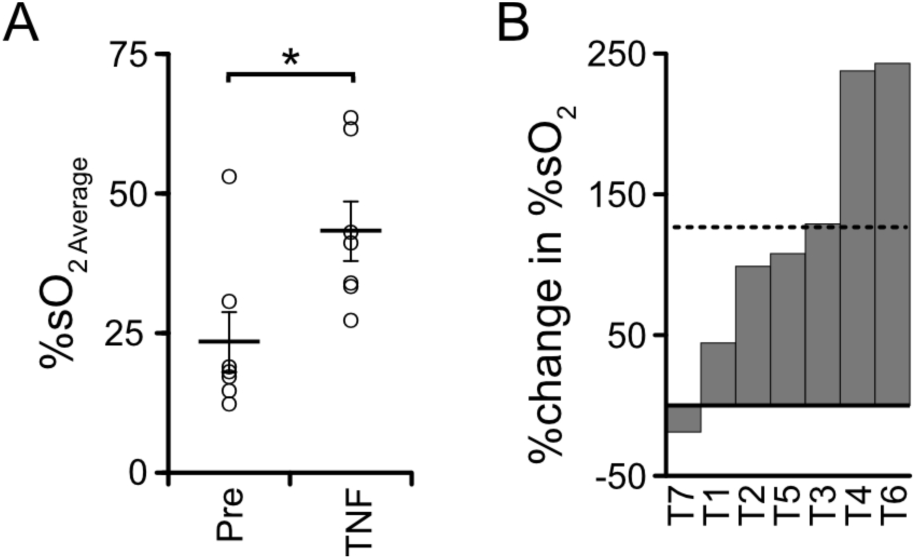
TNF in the absence of castration is insufficient to induce intra-tumoral hypoxia in Myc-CaP tumor. **A**, Mean PAI signal (%sO_2_) from tumors pre-castration (n=7), and at one day following TNF treatment (n=7). Mean (columns) and SEM (bars). *p<0.05. **B**, Waterfall plot of %change in %sO_2_ in individual tumors (columns) after TNF treatment. Mean %change is dashed line.

## Discussion

Previously, we reported that TNF, the prototypical death receptor ligand – but not other death receptor ligands such as FasL or TRAIL – was required for castration induced regression of the normal murine prostate (5). This is consistent with the original description of prostate cell death via apoptosis by Kerr *et al*. (3). Treatment of normal mice with sTNFR2-Fc did not completely block castration-induced regression, suggesting other apoptotic mechanisms are required for complete regression of the normal gland (5). In this report, we employed complementary functional imaging modalities and vessel structural analysis to confirm and extend the previous descriptions of a vascular component of regression. This pathological response to androgen deprivation – beginning with endothelial cell apoptosis and increased vessel permeability and culminating in hypoxia (12-15) – indirectly contributes to prostate cancer regression. In addition to a comprehensive analysis of vascular changes in the subcutaneous Myc-CaP model, we also detected castration-induced hypoxia (Fig. 5) in a genetically engineered PTEN-deficient prostate cancer model. We were not able to detect reduced blood flow by Power Doppler or reduced perfusion by CE-US in the PTEN-deficient model, perhaps because of limitations in imaging signal penetration for the endogenous prostate relative to the subcutaneous tumors in Figs. 2-4.

We demonstrated that micro-vessel damage using anti-CD31 immunohistochemistry, reduced blood flow via PD, reduced perfusion via CE-US and hypoxia via PAI, were all inhibited by blocking TNF signaling with sTNFR2-Fc. While we found a mixed micro-vessel density increase in tumors from different sTNFR2-Fc treated host animals after castration, we demonstrate TNF is necessary for castration-induced structural vessel damage, including diameter, area, and vessel wall thickness (Fig. 1). These events all occur at one day post-castration, prior to the onset of tumor shrinkage (Fig. S1). We have also recently observed that tumor volume in PTEN-deficient model does not decrease until three or more days following castration (34). It has been shown that apoptosis rates in the endothelium increase prior to epithelium (8), suggesting that endothelial damage occurs earliest. In addition, since we believe that the events in Figs. 1-4 are sequentially ordered (Fig 7) and since all are inhibited by sTNFR2-Fc, we conclude that TNF acts at the most proximal step. Indeed, there are multiple reports that TNF can induce both endothelial apoptosis and increase vascular permeability (16; 17; 35; 36).

**Figure 7.**
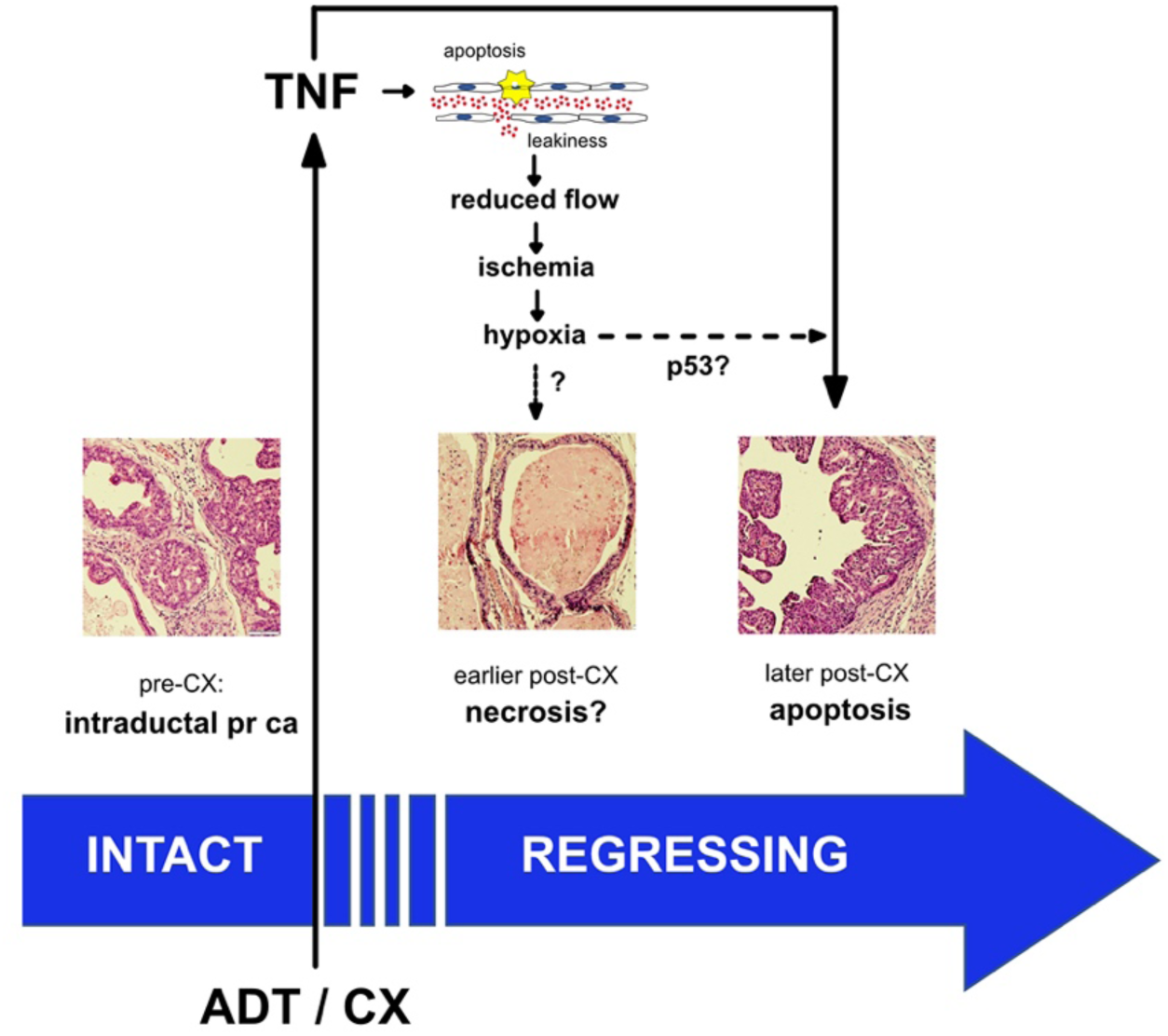
Proposed mechanism for TNF regulation of prostate regression. A proposed mechanism for prostate cancer regression following androgen deprivation therapy is illustrated, focusing on the contribution of events that are mediated by the tumor microvasculature. We propose that one of the earliest events is the activation of TNF signaling in endothelial cells leading to increased permeability (‘leakiness’) and endothelial cell death. The damage to the endothelium leads to a cascade of vascular changes – reduced blood flow, ischemia due to reduced perfusion and eventually transient tissue hypoxia – that likely enhances the death of the epithelial component of the tumor. A prediction of the model is that endothelial cell apoptosis precedes epithelial cell apoptosis, and this has been noted in the literature (see text). There is limited support for the usual cell death consequence of hypoxia (namely, necrosis or necroptosis), but there is a partial requirement for p53 in castration-induced regression, suggesting that p53-hypoxia signaling contributes to death receptor mediated apoptosis of the epithelial tumor cells.

Given that TNF signaling is the likely initiating signal for vascular regression, a key question is – what is the molecular trigger that activates TNF signaling in the endothelium? In the normal prostate, we observed an acute increase in TNF mRNA in the stromal compartment about 8 hours post-castration {Davis, 2011 #5243}. In addition, we and others previously showed that c-FLIP is down-regulated by castration via the androgen receptor acting on the c-FLIP promoter (37; 38). c-FLIP is a dominant-negative homologue of caspase-8 that acts as a natural inhibitor of death receptor signaling and its sustained expression may play a role in the development of castration resistant prostate cancer (39). In the Myc-CaP model investigated in this report, TNF levels do not change post-castration, suggesting that some other component of the TNF signaling cascade is regulated. Since the androgen receptor is not expressed in rodent endothelium (40) it seems unlikely that c-FLIP is a key regulator. Human endothelium, in contrast, do express AR and therefore the role of c-FLIP could be significant (7). Other components of the TNF apoptotic signaling network remain potential candidates for activating TNF signaling in the endothelium, post-castration.

Our results also suggest a second key question – how does hypoxia lead to epithelial cancer cell death? One obvious result of tissue hypoxia is necrosis (or perhaps necroptosis). We did detect necrosis in some tissue sections from the PTEN-deficient prostate cancer model (Fig. 7) but we have not been able to document this on a consistent basis, nor have we observed necrosis in the Myc-CaP model. PTEN-deficient prostate cancers have an intraductal histology, which is rarely seen in human prostate cancer but is possibly predisposed to hypoxia-induced necrosis, given the multiple layers of cancerous cells lining the ducts in these tumors. A more likely mechanism, especially in histologically typical prostate cancers, is that hypoxia activates or enhances epithelial apoptosis. Hypoxia-induced p53 (41) is a candidate inducer of apoptosis during prostate cancer regression. Most primary prostate cancers express wild type functional p53 (Fig. S2). Moreover, castration of p53 null mice yields partial regression of the normal prostate (42; 43), implying a role for wild type p53 in castration-induced regression. In the case of DNA damage stress, elevated p53 protein levels are well known to induce cell cycle arrest and apoptosis via caspase-9 and Bcl-2 family proteins. However, hypoxia induced p53 seems to function in a distinct fashion (44), with p53 transcriptionally regulating a distinct set of genes that encode apoptosis regulators (45). Therefore, p53 induction could be a key molecular signal for hypoxia induced epithelial apoptosis (Fig. 7). Our hypothesis also suggests that patients with mutant p53 prostate cancers will not respond as well to ADT.

Finally, the TNF-mediated vascular damage caused by ADT may provide an opportunity for prostate cancer therapy. TNF administration induces short-duration vascular disruption in many cancers, transiently enhancing tumor permeability and ultimately reducing blood flow (46-48). TNF therapy induces tumor regression in rodent models (49), but in humans results in hypotension and other dose-limiting toxicities, thus preventing effective use as a systemic anti-cancer therapy (50). Neovasculature-targeted TNF circumvents this toxicity by producing locally high TNF levels and induces vessel permeability (51), enhancing chemo- and immuno-therapy (52; 53). Similarly, tumor microenvironment targeted TNF enhances CD8+ T-cell mediated anti-tumor immunotherapy (54). We found that ADT, by inducing paracrine TNF signaling, disrupts the tumor vasculature in prostate tumors. However, the ADT-induced vascular damage is not durable, and tumor vascularization increased four weeks after castration (12). A similar transient effect is seen in human tumors (7). This suggests that ADT creates a window of vulnerability during which concurrently administered therapies may reduce cancer progression.

## Authors’ Contributions

<u>Conception and design</u>: M. Seshadri, J.J. Krolewski, and K.L. Nastiuk

<u>Development of methodology</u>: S. Singh, L.Rich, S. Turowski, K.L. Nastiuk

<u>Acquisition of data</u>: S. Singh, K. Sha, C. Pan

<u>Analysis and interpretation of data</u>: S. Singh, K. Sha, C. Pan, N. Jaiswal-Agrawal, L. Rich, J.J. Krolewski, and K.L. Nastiuk

<u>Writing, review, and/or revision of the manuscript</u>: J.J. Krolewski, M. Seshadri, K.L. Nastiuk

<u>Study supervision</u>: K.L. Nastiuk

## Acknowledgements

We thank the staff members of the Roswell Park Comprehensive Cancer Center’s Experimental Tumor Models, Translational Imaging, and Laboratory Animal Shared Resources for their assistance.

## Supplementary Figure

**Figure S1.**
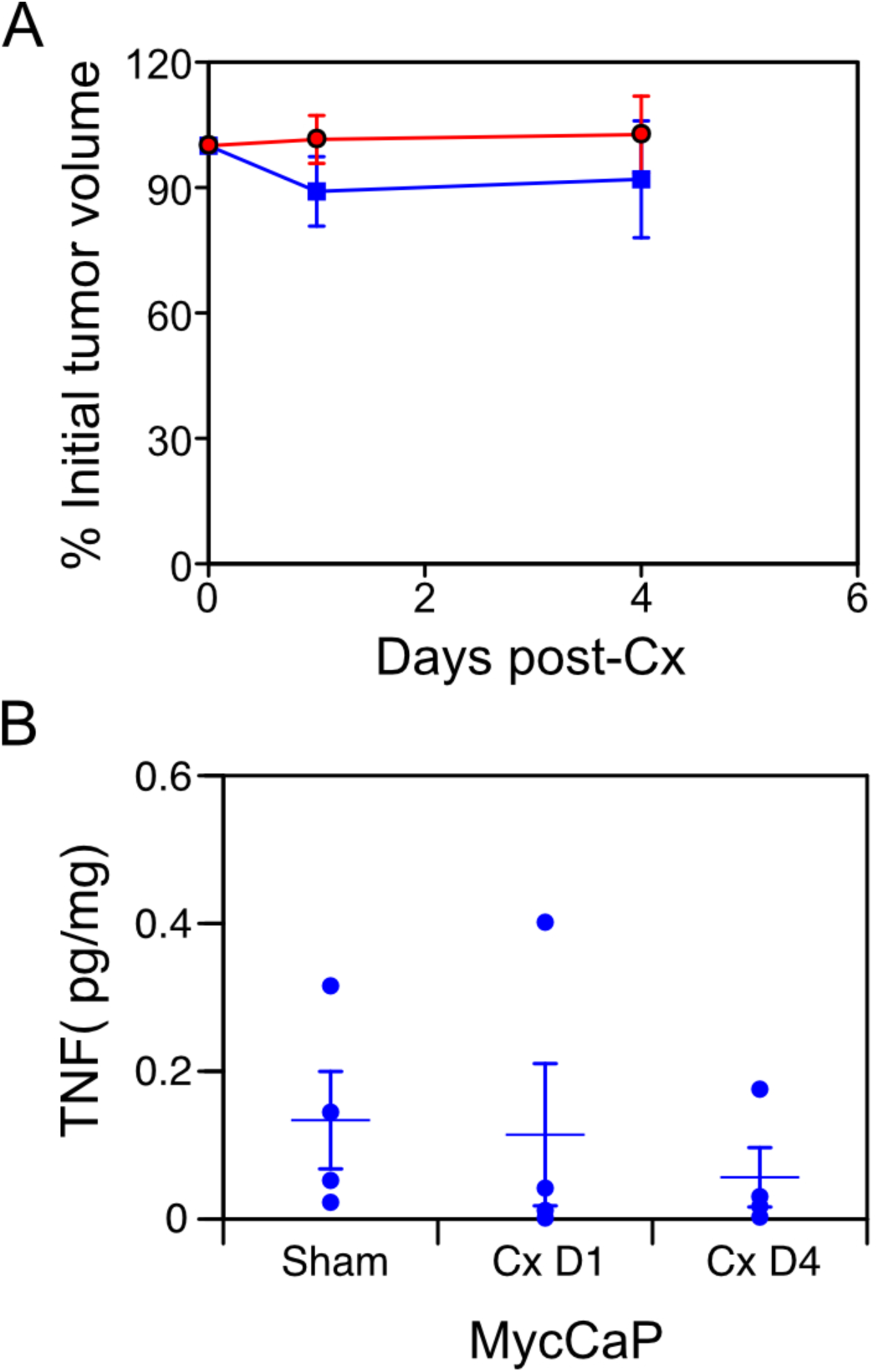
Tumor volume and TNF protein levels were not changed post-castration in prostate cancer allografts. **A**, HFUS determined Myc-CaP tumor volume after castration (normalized to pre-castration volume) in mice treated with vehicle (n=9, blue) or sTNFR2-Fc (n = 7, red). **B**, Tumor tissue was collected from FVB mice bearing Myc-CaP subcutaneous xenografts one and four days after castration (Cx) or from sham-castrated mice. TNF protein levels were measured in solublilized tumor tissue by ELISA (R&D #DY410), pg per mg soluble tumor extract, n = 4 tumors for each group, with individual tumor TNF quantities (circles), and group means (lines) and SEM (bars).

**Figure S2.**
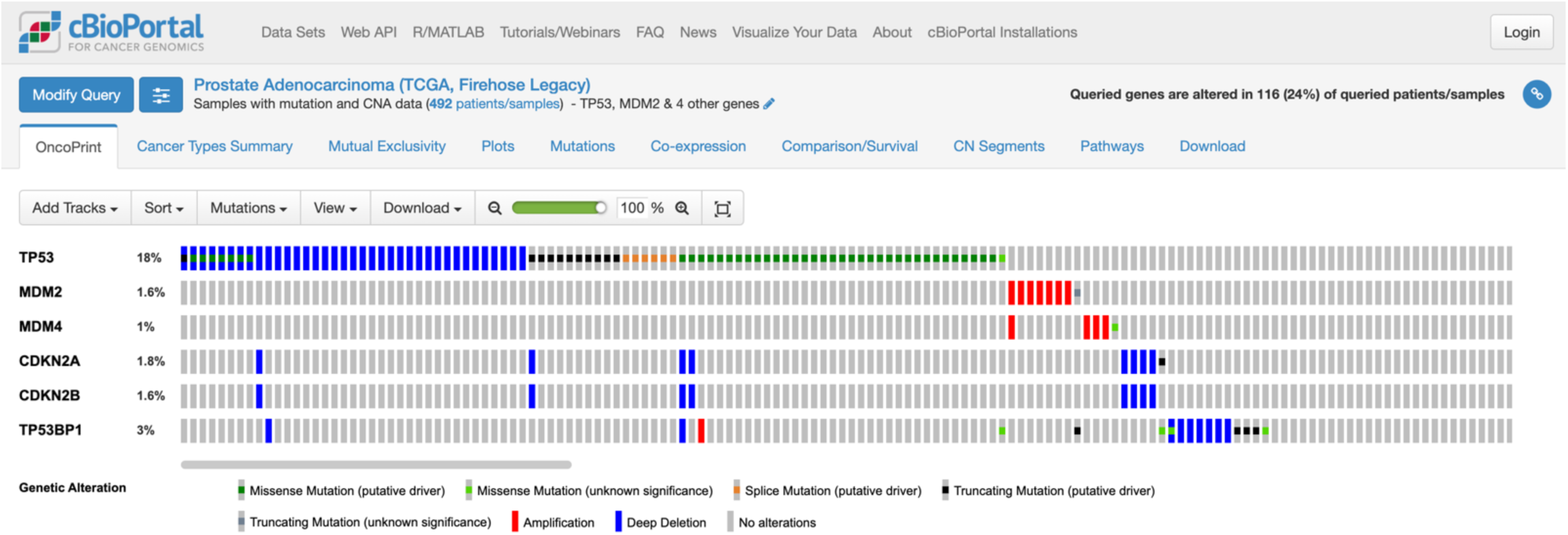
TP53 mutation frequency in primary prostate cancers. We used the cBioPortal web-browser to determine genomic alterations (mutations and copy number alterations) in the TCGA dataset indicated (TCGA, Firehouse Legacy) containing 492 primary prostate cancers. The key indicates the color-coding of the alterations. Grey bars have no alterations for the indicated set of 6 genes. Only a portion of the unaltered patient cases are shown.

